# Insights into diversity, host range, and evolution of iflaviruses in Lepidoptera through transcriptome mining

**DOI:** 10.1101/2025.01.22.634245

**Authors:** Devin van Valkengoed, Astrid Bryon, Vera I.D. Ros, Anne Kupczok

**Affiliations:** Bioinformatics Group, Wageningen University, 6700 PB Wageningen, The Netherlands; Princess Máxima Center for Pediatric Oncology, 3584 CS Utrecht, The Netherlands; Laboratory of Virology, Wageningen University, 6700 PB Wageningen, The Netherlands

## Abstract

Insects are associated with a wide variety of diverse RNA viruses, including iflaviruses, a group of positive stranded RNA viruses that mainly infect arthropods. Whereas some iflaviruses cause severe diseases in insects, numerous iflaviruses that were detected in healthy populations of butterflies and moths (order: Lepidoptera) do not show apparent symptoms. Compared to other hosts, only few iflavirus genomes for lepidopteran hosts could be found in publicly available databases and we know little about the occurrence of iflaviruses in natural and laboratory lepidopteran populations. To expand the known diversity of iflaviruses in Lepidoptera, we developed a pipeline to automatically reconstruct virus genomes from public transcriptome data. We reconstructed 1,548 virus genomes from 55 different lepidopteran species that we were identified as coding-complete based on their length. To include incompletely assembled genomes, we developed a reference-based patching approach, resulting in 240 patched genomes. By including publicly available genomes, we inferred a phylogeny consisting of 139 non-redundant iflavirus genomes. Of these, 65 represent novel complete genomes, of which 39 might even belong to novel virus species. Our analysis expanded virus host range, where highly similar viruses were found in the transcriptomes of different lepidopteran species, genera or even families. Additionally, we find two groups of lepidopteran species depending on the diversity of viruses that infect them: some species were only infected by closely related viruses, whereas other species are infected by highly diverse viruses from different regions of the phylogeny. Finally, we show that the evolution of one virus species, *Iflavirus betaspexiguae*, is impacted by recombination within the species, which is also supported by the co-occurrence of multiple strains within data sets. Our analysis demonstrates how data mining of publicly available sequencing data can be used at a large scale to reconstruct intra-family viral diversity which serves as a basis to study virus host range and evolution. Our results contain numerous novel viruses and novel virus-host associations, including viruses for relevant insect pests, highlighting the impact of iflaviruses in insect ecology and as potential biological control agents in the future.

## Introduction

RNA viruses are abundantly present in arthropods (Wu et al. 2020; Qi et al. 2023). Within the order *Picornavirales*, the *Iflaviridae* family contains only one genus (*Iflavirus*) and these viruses mainly infect insects, spiders, mites and crustaceans. Iflavirus infections can range from asymptomatic to severe, causing abnormalities or death in natural insect populations and in mass rearings of economically interesting insects (Valles et al. 2017). Furthermore, it is shown that the most common route of infection among iflaviruses is through horizontal transmission by ingestion of virus-contaminated food sources, but also vertical transmission has been reported (Amiri et al. 2016; van Oers 2010). Transmitted both horizontally and vertically, sacbrood virus and deformed wing virus in honey bees cause disastrous infections leading to diarrhea and developmental malformations in larvae and pupae, and morphological abnormalities in adults, respectively (de Miranda & Genersch 2010; Wei et al. 2022; Yue et al. 2007). Besides the iflaviruses detected in honey bees and ants (order: Hymenoptera), several members of this viral family have been detected in other arthropods like brown planthoppers and aphids (order: Hemiptera), flies and mosquitoes (order: Diptera), Varroa mites (*Varroa destructor*; order: Mesostigmata), ticks (order: Ixodida), wild and cultivated house crickets (order: Orthoptera), and butterflies and moths (order: Lepidoptera) (Murakami et al. 2014; Jiao et al. 2024; Meki et al. 2021; Guimarães et al. 2024; Ongus et al. 2004; Daveu et al. 2021; de Miranda et al. 2021).

In lepidopteran insects, some iflaviruses do not show clear symptoms and have been discovered in apparently healthy insect populations (van Oers 2010), like Spodoptera exigua iflavirus 1 (SeIV1) (Millán-Leiva et al. 2012) and Spodoptera exigua iflavirus 2 (SeIV2) (Choi et al. 2012; Jakubowska et al. 2014) detected in the beet armyworm *Spodoptera exigua* (family: Noctuidae). The high prevalence of both iflaviruses in field and laboratory populations suggests that they are of ecological importance in *S. exigua* (Jakubowska et al. 2014; Virto et al. 2014). While effects of SelV2 are unknown, SeIV1 is shown to have minor fitness effects on its host (Carballo et al. 2020). In other lepidopteran species, iflaviruses cause severe symptoms, including mortality. For example, tremendous cocoon losses in silk farms are caused by the *Bombyx mori*-infecting infectious flacherie virus (IFV), which was the first iflavirus for which a complete genome sequence became available (Isawa et al. 1998).

Iflaviruses form non-enveloped icosahedral particles that are between 22 and 30 nm in diameter (Valles et al. 2017). Members of the *Iflaviridae* family have a positive sense, single-stranded RNA (+ssRNA) genome, which is non-segmented and varies between 9 and 11 kb in size. The genome is led by a 5’ untranslated region (UTR) including the presence of an internal ribosomal entry site (IRES) for some iflaviruses and harbors a 3’ UTR followed by a poly-A tail. Iflavirus genomes encode a single open reading frame (ORF) that translates into a polyprotein of ca. 3000 amino acids, which is post-translationally cleaved into functional proteins (van Oers 2010). At the 5’ end, preceded by a leader protein, the structural capsid proteins are located, followed downstream by the non-structural proteins (van Oers 2010). The International Committee on Taxonomy of Viruses (ICTV) currently recognizes 16 iflavirus species. Phylogenetic analysis showed that iflaviruses group into different clades and no obvious evolutionary relationship based on insect host is noted, since each iflavirus clade is composed of viruses that infect different insect orders.

RNA viruses are known to evolve rapidly and to have high mutation rates (Duffy 2018; Butković & González 2022). Additionally, especially +ssRNA viruses are known to recombine frequently, which might be linked to template switching of the RNA polymerase, where co-infection of the same cell is required for recombination to happen between two viruses (Butković & González 2022; Wells et al. 2023; Pérez-Losada et al. 2015). Since some nucleotide similarity is required for template switching, recombination typically happens between related virus variants; nevertheless, recombination between more diverged variants can also occur (Simon-Loriere & Holmes 2011).

Many studies in virology traditionally focus on human-infecting viruses, by sequencing human samples or vectors transmitting these viruses (e.g., mosquitos, Pan et al. 2024). Recently, research expanded to sequence-based virus detection from diverse biological samples, including non-model organisms. This approach led to a tremendous increase in the known RNA virus diversity over the past few years that was pioneered by virus detection in invertebrate metatranscriptomes (Shi et al. 2016). This advance was fueled by decreasing costs and increasing amounts of sequencing data and by improved algorithms for virus detection. The methods used comprise approaches that detect RNA-dependent RNA polymerase (RdRp) based on sequence search (e.g., RdRp-scan, Charon et al. 2022) or artificial intelligence (Hou et al. 2024). Recently, virus discovery is also powered by the re-use of existing data, also called data mining, which is cost-efficient and allows to cover species where it can be difficult to obtain samples, such as protected species (Harvey et al. 2023). For example, the search for relatives of human pathogens in animal datasets allows to study their origin and evolution, as exemplified by studies on Hepatitis B virus (Lauber et al. 2017), Hepatitis delta virus (Iwamoto et al. 2021), and coronaviruses (Lauber et al. 2024). Relatives of insect-transmitted human viruses, such as flaviviruses, were also detected in diverse animals (Mifsud et al. 2023; Paraskevopoulou et al. 2021). Next to this, data mining substantially enhanced the knowledge of virus diversity in fungi (Gilbert et al. 2019), plants (Sidharthan et al. 2024; Mifsud et al. 2022; Bejerman & Debat 2022), and insects (Wu et al. 2020). Additionally, metatranscriptome analysis of environmental communities expanded the known diversity of bacteriophages and eukaryotic viruses (Neri et al. 2022; Zayed et al. 2022).

For known virus families, typically only few complete genomes have been deposited in databases, whereas many more could be found in sequencing data, such as transcriptome assemblies (Olendraite et al. 2023). The number of novel viruses that can be found in public data is especially high for non-vertebrate hosts and those novel viruses include many iflaviruses (Olendraite et al. 2023). Potentially even more viruses can be detected in sequencing reads that are deposited in the sequence read archive (SRA), as shown by the Serratus platform that provides an inventory of SRA data sets showing RdRp signatures of particular virus families (Edgar et al. 2022). Despite the recent massive increase in known RNA viruses, we still know little about virus diversity and evolution (Harvey & Holmes 2022). Expanding the genome diversity of known virus families would enable in-depth investigations of the sequence evolution of these viruses. As only 27 iflavirus genomes for lepidopteran hosts could be found in the GenBank database at the National Center for Biotechnology Information (NCBI), in contrast to 266 for other hosts, we here aim to reconstruct iflavirus genomes from public SRA libraries of diverse lepidopteran species, which provides a basis to study iflavirus diversity, host range, and evolution.

## Results and Discussion

### Most detected *Picornavirales* in Lepidoptera belong to *Iflaviridae*

To study the diversity of *Picornavirales* in Lepidoptera, we used the Serratus database to obtain SRA libraries with viral signals, which we used as input for our analysis. We developed a pipeline to map the libraries to the insect host genome (if available), assemble the unmapped reads, and detect contigs with sequence similarity to viruses (Figure 1A). In total, the assemblies of 4,002 Lepidoptera RNA-Seq SRA libraries were checked for sequences belonging to *Picornavirales*. For *Iflaviridae*, we termed a genome as complete if it has a match to an *Iflaviridae* protein and is between 8 and 12kb in length. We found in total 1,548 complete contigs that contained an ORF of at least 6,000 nt from 1,483 different libraries, where an accession could contain up to 3 complete genomes (Supplementary tables: Table S1, Figure 1B). In addition, if the assembly contains multiple hits to *Iflaviridae* proteins that are shorter than 8kb, this genome is considered fragmented (384 fragmented, Supplementary tables: Table S1). Of these, we could patch 240 to a near-complete polyprotein (see Methods). Thus, from the 3,214 libraries, where Serratus showed signals of *Iflaviridae*, we could reconstruct complete or fragmented genomes for 1,709 of these (53%, Supplementary tables: Table S1). In addition, we included 15 libraries from *S. exigua* that were not detected with Serratus. These have been included nonetheless because they belong to experiments where iflaviruses have been detected suggesting that they might also contain viral reads. Of these, complete genomes could be reconstructed from seven libraries and a patched genome from one library, showing that Serratus can miss virus-containing libraries.

**Figure 1.**
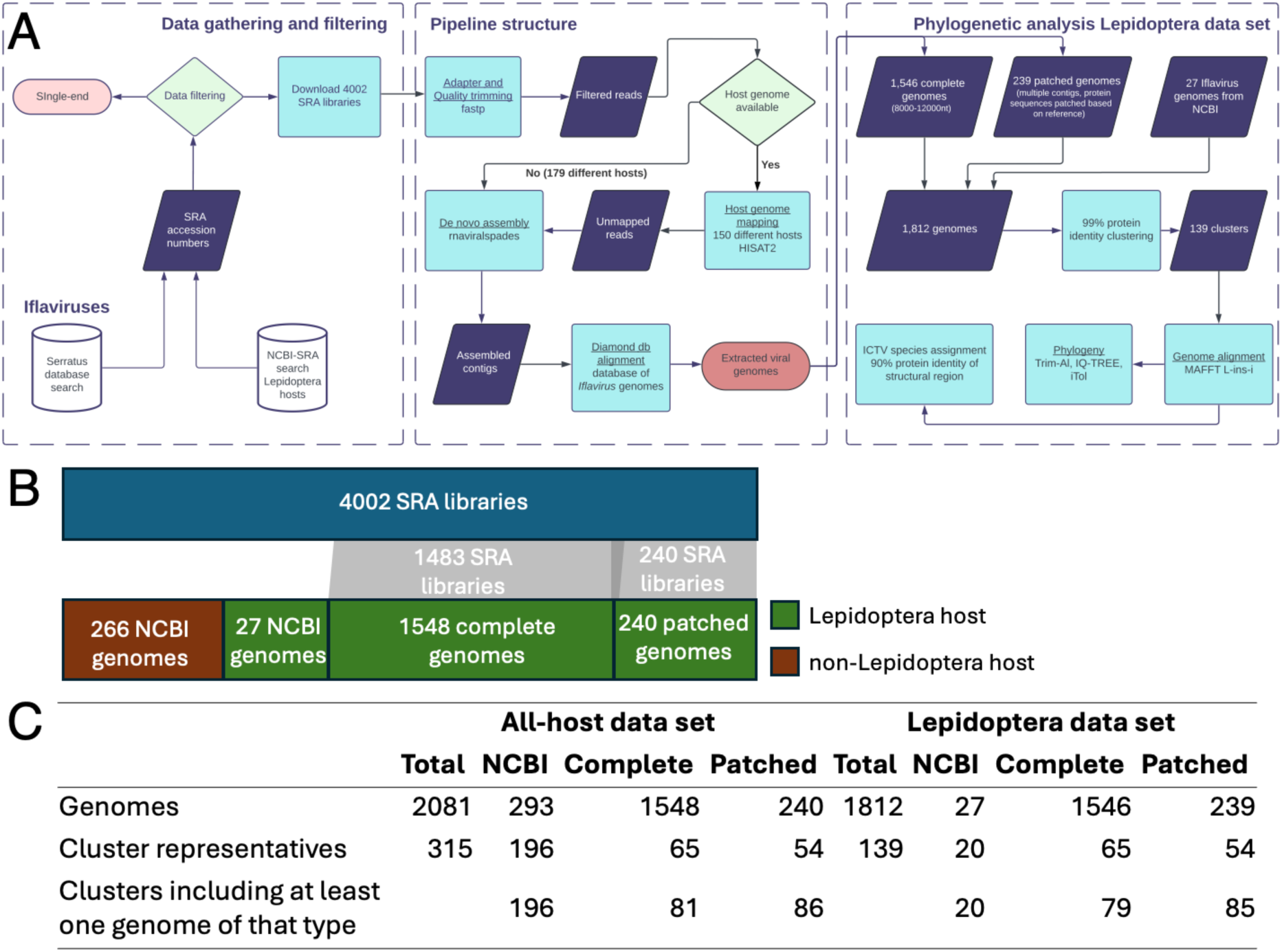
Workflow and data set overview. (A) Computational workflow. The phylogenetic analysis provides the numbers for the Lepidoptera data set. (B) Numbers of processed libraries and included genomes. (C) Overview of included genomes in both data sets. There are 2 complete and 1 patched genome missing from the Lepidoptera data set since they are in a cluster with a non-Lepidoptera representative: SRR11614424_complete_1 from host *Parnara guttata* clusters with UHR49772.1 from host *Tetragnatha nitens*, SRR9171794_complete_1 from host *Heliozelidae sp.* clusters with YP_009305421.1 (Moka virus) from host *Vespula pensylvanica*, ERR10123688_pg_nr1 from host *Synanthedon andrenaeformis* clusters with AUK50716.1 (Sacbrood virus) from host *Apis mellifera*.

In addition, we found 479 contigs with hits to *Picornavirales* genomes that belong to other families than *Iflaviridae* or that could not be classified at family level (Supplementary tables: Table S1). To conclude that these are actual viral contigs and to assess their completeness, these contigs would need to be further analyzed and filtered to include only long alignments. Since iflaviruses are found at much higher numbers compared to the other families within the *Picornavirales*, we restricted our subsequent analyses to the family of *Iflaviridae*. The sequences with similarity to other *Picornavirales* and all the complete and patched iflavirus genomes can be found in GitHub (https://github.com/annecmg/VirusRePublic/tree/main/Data_Paper_Lepidoptera).

The 4,002 analyzed libraries originate from 307 different lepidopteran species. We notice that many of the reconstructed iflaviruses were found in well studied lepidopteran species, for example in *Bombyx mori* and *Spodoptera litura* (Supplementary tables: Table S3, Figure S1). For some lepidopteran species, we reconstructed a large number of iflavirus genomes. For these lepidopteran species, often multiple libraries of one BioProject were present, suggesting that the same insect population was studied under different conditions in a transcriptomic experiment. Although Serratus detected signals of *Iflaviridae* in libraries from 217 lepidopteran species, we only reconstructed iflavirus genomes from 70 lepidopteran species. For some of the lepidopteran species that lack reconstructed iflavirus genomes, many libraries were detected (e.g., 82 libraries from one BioProject from *Melitaea cinxia* or 75 libraries from four different BioProjects from *Papilio xuthus*, Supplementary tables: Table S3).

Taken together, we could reconstruct complete iflavirus genomes for about half of the samples where virus signatures were detected (Supplementary tables: Tables S1, S3). This might be partly due to the loose settings used for Serratus, which can lead to false-positive predictions. Alternatively, our assembly-based approach could also miss viruses, particularly when viruses are of low abundance or are highly diverse. Another possible explanation, that might be applicable to the lepidopteran species where no iflavirus genomes could be reconstructed, is that our filtering criteria missed iflaviruses without sequence similarity to known iflaviruses or with a different genome architecture.

### Many reconstructed iflavirus genomes have similarity to known iflavirus genomes

To reduce redundancy among the 2,081 iflavirus genomes (1,548 complete, 240 patched, 293 from NCBI), we clustered them based on pairwise blast (99% identity, 95% alignment coverage), resulting in 93 clusters (each with 2 to 307 sequences) and 222 singletons (Figure 1C). When assigning cluster representatives, we preferred NCBI genomes over complete genomes over patched genomes. Of the 315 cluster representatives, only 139 have lepidopteran hosts. In the following, we denote the full data set of 2,081 genomes and 315 clusters that includes all NCBI iflavirus genomes as “All-host data set”. In contrast, the “Lepidoptera data set” includes only the clusters whose representatives have a lepidopteran host, resulting in 1,812 genomes and 139 clusters in total. Of the 139 clusters in the Lepidoptera data set, only 20 include a genome from NCBI, whereas 65 additional clusters include a complete genome (Figure 1B). In total, 536 (35%) complete genomes fall into 14 different clusters with an NCBI genome, whereas the remaining ones show less than 99% amino acid identity to a known NCBI genome. Thus, we found that a substantial proportion of the reconstructed iflavirus genomes shows high similarity to known iflavirus genomes, supporting the accuracy of our approach. Beyond that, we also reconstructed many viral genomes that are divergent to the known iflavirus genomes. The 65 cluster representatives originating from the reconstructed complete genomes are named strain WUR1 to WUR65 and can be found at NCBI (BioProject PRJNA1137245). All reconstructed iflavirus genomes can be found in GitHub (https://github.com/annecmg/VirusRePublic/tree/main/Data_Paper_Lepidoptera).

### The majority of the reconstructed complete iflavirus genomes is coding-complete

As the genome completeness criteria were only based on protein length, the set of reconstructed iflavirus genomes we termed complete could still include incomplete genomes. To assess the genome completeness, we compared the assembly statistics of the complete genomes with that of the NCBI genomes. We observed that the reconstructed genomes tend to be longer than the NCBI genomes and that the UTRs are comparable in size to the NCBI genomes (Figure S2). A 3’ UTR of length less than 3nt indicates that the open reading frame (ORF) stopped because the contig ended; thus likely no stop codon was included and the ORF is incomplete at the 3’ end. Of all complete genomes, 47 (3.0%) have an incomplete 3’ UTR, suggesting incomplete coding sequences. However, when only considering cluster representatives, only one genome (1.5%) has an incomplete 3’ UTR (WUR9 SRR3089402_complete_1).

Additionally, we annotated the cluster representatives with InterPro to compare the domain content of the reconstructed genomes to that of the NCBI genomes (Supplementary tables: Table S4). All the domains that are observed in NCBI genomes from the Lepidoptera data set can also be observed in the complete genomes. Furthermore, our reconstructed genomes do not contain any additional domains. The domains found in 19 of the 20 NCBI genomes from the Lepidoptera data set are also found in the complete genomes, with one exception: WUR9, which is identified above as having an incomplete 3’ UTR, lacks the polymerase which is located at the C-terminus of the polyprotein due to a truncation at the C-terminus.

We thus conclude that most of the reconstructed genomes that are termed complete here have complete coding sequences, and this holds for nearly all of the reconstructed representatives. Note that our definition of completeness is only based on the contig length and matches to known iflavirus genomes; however, we did not evaluate whether the genomes are indeed complete in a strict sense. To assess the completeness of the UTRs, further analyses would be needed; for example, mapping the reads to contigs could confirm if there is a clear drop off at the end as expected for nucleotide-complete sequences. Coding-complete sequences are adequate to use for our analyses that focus solely on coding sequences. In agreement with previous studies, we show that coding-complete virus genomes can successfully be recovered from transcriptomics data (e.g., Lauber et al. 2024).

### We find known and novel genomes throughout the iflavirus phylogeny

To infer how the genomes reconstructed here are related to known iflavirus genomes, we reconstructed a phylogeny of the 315 cluster representatives from the all-host data set. We observed that the reconstructed genomes can be found in different parts of the iflavirus phylogeny and that most of them group with other genomes found in Lepidoptera (Figure S3). Nevertheless, some reconstructed genomes group with viruses found in other orders than Lepidoptera; these relationships typically show long branches in the phylogeny, i.e., the reconstructed viruses are only divergently related to the non-Lepidoptera viruses. For example, WUR48-WUR52 form a divergent group of viruses that is found in four different lepidopteran species belonging to different families (*Eogystia hippophaecolus, Cydia nigricana, Heliozelidae sp., Parnassius cephalus*), grouping with *Lysiphlebus fabarum* RNA-virus type A and B found in Hymenoptera. There is only one instance in the phylogeny where reconstructed genomes are closely related to viruses found in non-Lepidoptera: three patched genomes have been reconstructed from transcriptomes of the pine pest *Dendrolimus punctatus*. The assembled genomes originate from the same project (PRJNA590087) and group closely with *Apis melifera* deformed wing virus. Due to the high similarity of the viral genomes from the *D. punctatus* libraries with the deformed wing virus, it is possible that the *D. punctatus* samples have been contaminated or the moths might have been exposed to the bee virus in the shared environment. The latter has been observed before for deformed wing virus and for other insects that share the environment with managed honey bees (Levitt et al. 2013). Similar observations can be made for the three reconstructed genomes that can be found in clusters together with viruses from other hosts than Lepidoptera; these could also be explained by contamination or ecological interactions (Figure 1C).

In the following, we focus on the Lepidoptera data set with 139 representative genomes. The maximum likelihood phylogeny is well supported with generally high bootstrap values (Figure 2). To infer to which iflavirus species the reconstructed genomes belong, we assigned virus species to the complete genomes according to the ICTV criteria, i.e., the amino acid sequence identity of the capsid region is at least 90%. We reconstructed complete genomes for all seven iflavirus species that are known to infect Lepidoptera (Supplementary tables: Table S5). Thereby, we substantially enhanced the genomic information and host range available for these known iflavirus species. For example, the host range for *Iflavirus spexiguae*, which includes *Spodoptera exigua* iflavirus 1, was extended to *Spodoptera frugiperda*, a species belonging to the same genus as *S. exigua* and both insects are known as significant agricultural pests. Furthermore, complete genomes of *Iflavirus betaspexiguae*, which includes *Spodoptera exigua* iflavirus 2, were also detected in lepidopteran species belonging to different families, such as *Spodoptera littoralis, S. litura, Lymantria dispar, Acrolepiopsis sapporensis* and *Galleria mellonella* which were previously not described as hosts of *Iflavirus betaspexiguae.* Some iflaviruses are known to cause lethal infections in a particular species, for example, *Iflavirus ectobliquae* in the tea looper (*Ectropis obliqua*) (Lu et al. 2007; Wang et al. 2004). Here, we also obtained a complete *I. ectobliquae* genome in another major tea plant defoliator, the tea geometrid *Ectropis grisescens* (Li et al. 2019). These findings highlight the potential of this iflavirus as a biological control agent against these tea plant pests. Another economically relevant virus is the *Antheraea pernyi iflavirus* (*I. vomitus*), which is indicated as the causative agent of the *Antheraea pernyi* vomit disease in the Chinese oak silkworm (*A. pernyi),* an important silk producing species (Geng et al. 2017). Complete genomes of this iflavirus were also assembled from libraries of the wild silk moth (*Bombyx mandarina)* and the Japanese silk moth (*Antheraea yamami),* suggesting a potential threat to other silk-producing lepidopteran species. Another noteworthy observation is made for *Iflavirus pernudae,* which is described to infect the host *Perina nuda*, a major defoliator of forest and shade trees in Asia (Wu et al. 2002; Cheanban et al. 2017). We also assembled complete genomes of this virus from libraries of *S. frugiperda, S. litura* and *B. mori* insect cells. To our knowledge, *I. pernudae* is not yet described in the aforementioned lepidopteran species and therefore we suspect that this virus is artificially induced, replicated and maintained in these specific cell lines.

**Figure 2:**
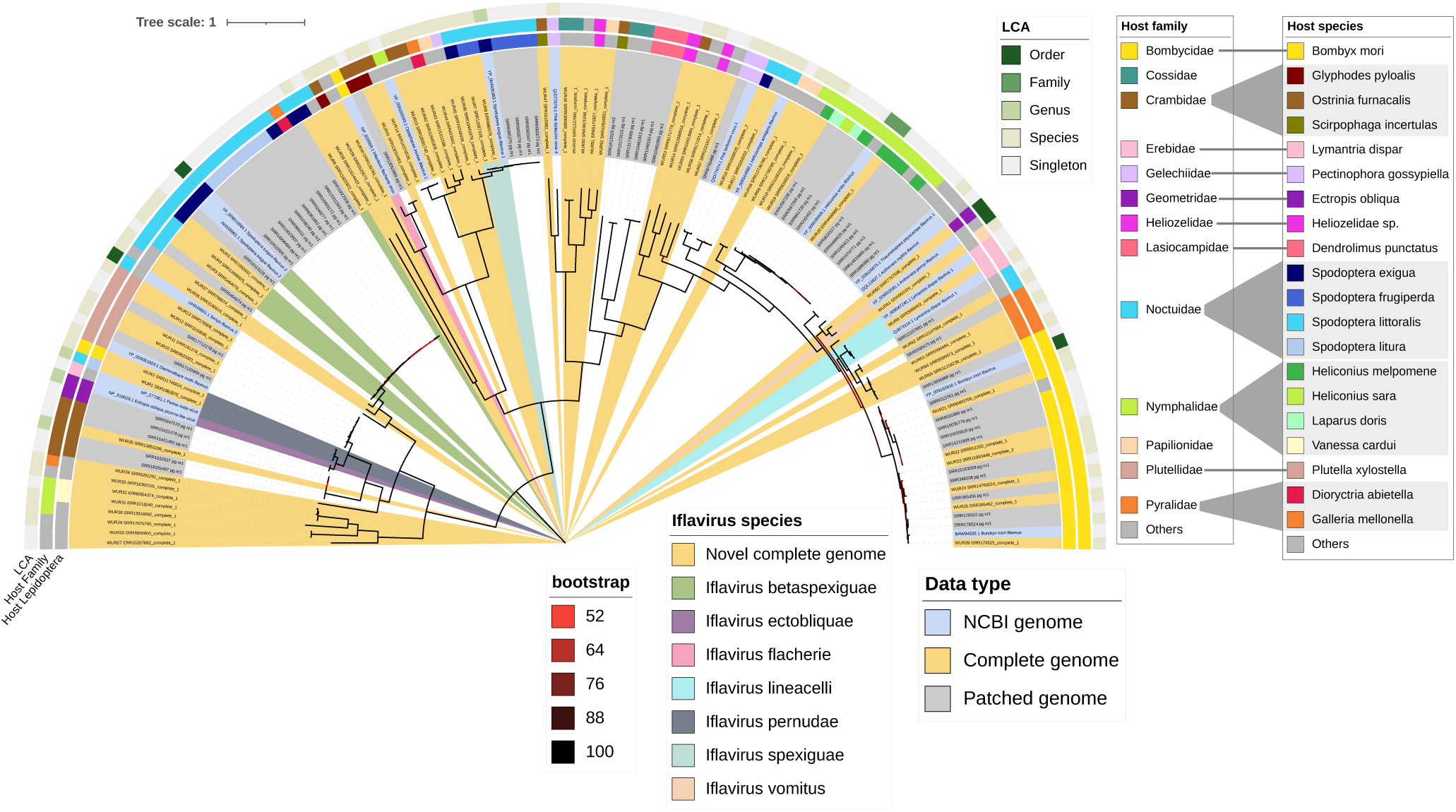
Iflavirus phylogeny for Lepidoptera data set. The Lepidoptera host species and family is shown in the outer ring; note that the host species is only given for the cluster representative, only host species and families occurring in at least three clusters are colored. Taxa name is colored by data type, clades are colored by iflavirus species, and branches are colored by bootstrap value. A completely annotated phylogeny can be found at https://itol.embl.de/tree/312010331252431721380571.

In the previous section, we have described that the diversity and host range within iflavirus species have been expanded in our analysis. However, not all complete genomes were assigned to a species recognized by ICTV. These remaining genomes fall into two different categories depending on whether a genome related at species level according to ICTV exists at NCBI or not (i.e., sequence identity of the capsid region of at least 90%). For the first category, a genome related at species level exists in NCBI, however, the related genome has also not yet been assigned to a viral species. These 17 cluster representatives thus belong to known iflaviruses and are marked white for “Iflavirus species” in Figure 2. One example of a virus that has not yet been assigned to a species is *Heliconius erato* iflavirus (Smith et al. 2014). We found multiple variants of this virus in seven different species of the Nymphalidae family (five *Heliconius* species, *Eueides isabella*, *Laparus doris*), indicating that this virus is abundantly present in different lepidopteran families.

The second category encompasses cluster representatives for which no related NCBI genome could be identified at the species level. These 39 complete genomes putatively belong to “novel” species and are thus marked “novel complete genome” in Figure 2. We observed reconstructed novel genomes throughout the iflavirus phylogeny (Figure 2), and they were detected in lepidopteran species with varying ecological impact. We find novel iflaviruses in Lepidoptera where some iflaviruses are known already: for example, WUR60 is found in *Ectropis obliqua*, host of *Iflavirus ectobliquae*, and WUR46 is found in *Spodoptera exigua*, host of *Iflavirus spexiguae* and *Iflavirus betaspexiguae*. In addition, we found novel iflaviruses in 23 different insect species, where no iflavirus has been described before. Some of these are major pest species like the bagworm, *Metisa plana* (WUR27, WUR28, WUR29), the tomato leafminer, *Tuta absoluta* (WUR45), the honeycomb moth, *Galleria mellonella* (WUR63, WUR64, WUR65) and the Asian corn borer, *Ostrina furnacalis* (WUR35). Furthermore, WUR30 is found in different geographic locations in the ecologically relevant butterfly species; the squinting bush brown (*Bicyclus anynana*) (Murugesan et al. 2022). For some of the novel viruses with complete genomes, it appears that they can infect multiple lepidopteran hosts. For example, WUR60 is found in the tea pest *Ectropis obliqua* and in *Dioryctria abietella*, a moth feeding on conifers. For some lepidopteran species, we even reconstructed multiple novel genomes of distantly related iflaviruses: *Dioryctria abietella* (WUR43, WUR60), the mulberry pest *Glyphodes pyloalis* (WUR39, WUR41), and the butterfly *Papilio polytes* (WUR44, WUR61), pinpointing to an understudied virome in some lepidopteran species.

### Iflaviruses can have a wide host range and highly diverse iflaviruses can be found in the same host

In the previous section, we highlighted examples of viruses from the same species infecting different hosts. This raises the question of how similar the host range of closely related viruses is. Note that the host given in Figure 2 is inferred only from the cluster representative. To learn whether this host is characteristic for the whole cluster, we reconstructed the lowest common ancestor (LCA) of all hosts in a viral cluster. Of the 62 non-singleton clusters, 51 (82%) contain only viruses from the same lepidopteran species, whereas the remaining eleven viruses infect different species of which five were even detected in different lepidopteran families (Figure 3A, Figure 2). For example, WUR20 from *Heliconius melpomene* was also assembled from libraries of other members of the Nymphalidae family like the Julia butterfly (*Dryas iulia)* from the Americas and the widespread painted lady (*Vanessa cardui*).

**Figure 3:**
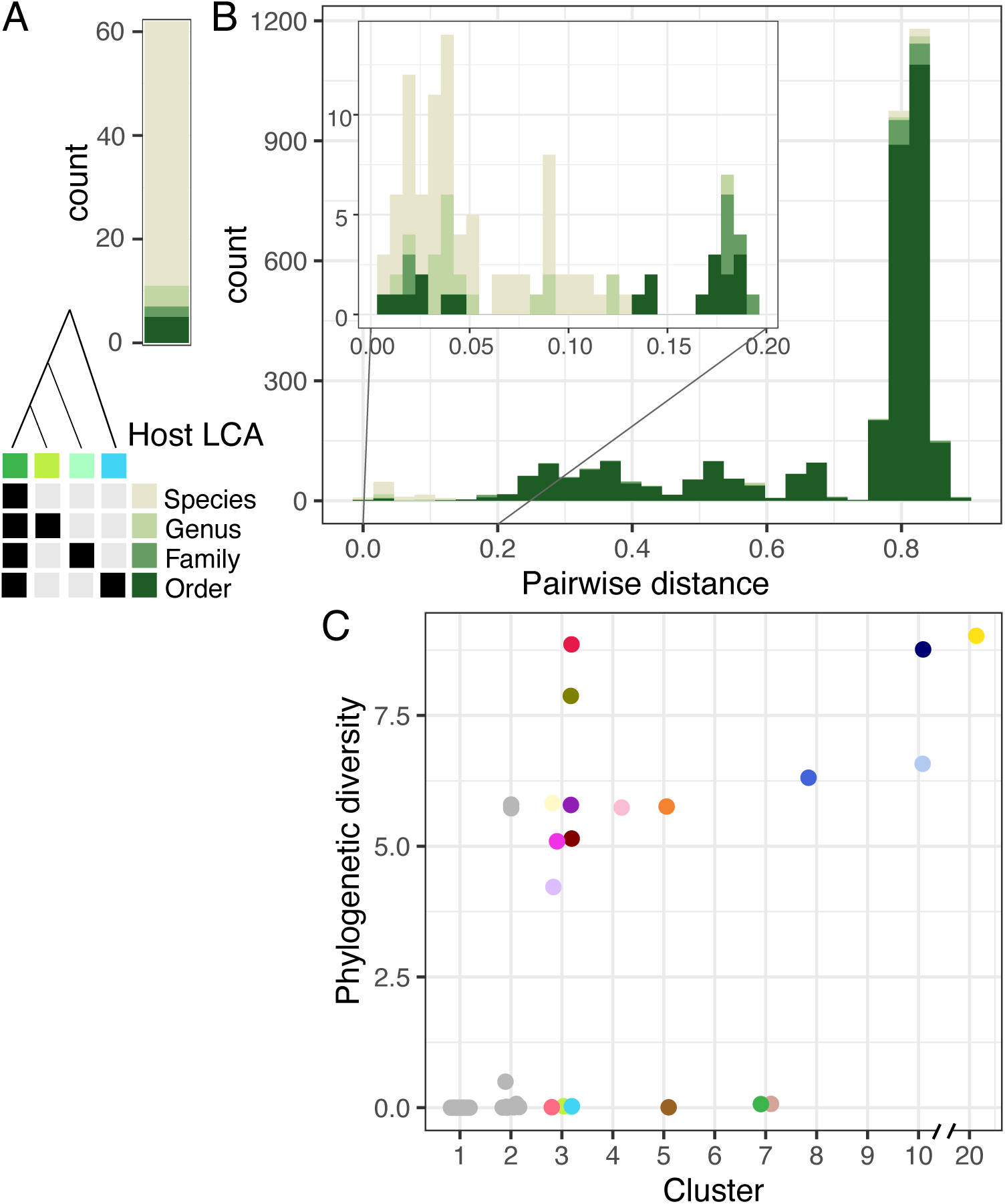
Host range. Lowest common ancestor (LCA) of Lepidoptera hosts (A) within all members of a cluster (also marked as LCA in Figure 2) and (B) for pairs across all cluster representatives, excluding patched genomes. Pairwise distance is given by 1 – amino acid sequence identity of the capsid proteins. The legend provides examples for the LCA of four different species (see color legend in Figure 2) based on their presence and absence in a pair. (C) Phylogenetic diversity (PD) of viruses infecting a particular Lepidoptera species (colored by host species according to Figure 2).

When comparing pairs of cluster representatives, we find that similar pairs (i.e., amino acid distance of the capsid region up to ∼13%) generally have the same host species (Figure 3B). Nevertheless, highly similar iflavirus pairs can also be found in different host species, genera or families. For example, we detected viruses that are highly similar to *Heliconius erato iflavirus* in other *Heliconius* species (WUR18, WUR19, WUR20). Highly similar viruses were also found across different lepidopteran genera, for example, WUR16 (assembled from *S. exigua* libraries) and *Helicoverpa armigera iflavirus* (initially described in host *H. armigera)*. An example of highly similar iflaviruses found in different host families is the *Opsiphanes invirae* iflavirus infecting *Opsiphanes invirae (*family Nymphalidae). We found that this iflavirus is highly similar to WUR14 and WUR15 assembled from libraries of the sugarcane borer *Diatraea saccharalis* (family Crambidae), which is a serious pest due to larvae boring into the sugarcane stalks.

Furthermore, virus pairs of cluster representatives with a distance higher than 20% generally have lepidopteran hosts that belong to different families (Figure 3B, pairwise distance >0.2 and LCA Order). However, we also observed that diverse iflaviruses can infect one lepidopteran species; especially for highly dissimilar viruses (Figure 3B, pairwise distance > 0.75 and LCA Species). To quantify the diversity of viruses that can be found in one insect species, we calculated the phylogenetic diversity (PD) of the viruses found in a particular species (Figure 3C, Supplementary tables: Table S3). We observed that for 46 species, only closely related viruses are known (up to a PD of 0.5), whereas 15 species can be infected by diverse viruses (with a PD larger than 4). Viruses from four well-investigated lepidopteran species, *B. mori*, *S. exigua, S. litura*, and *S. frugiperda*, were found in at least eight virus clusters and have a high phylogenetic diversity. In addition, for twelve lepidopteran species, we detected less than six virus clusters that represent a diverse group of iflaviruses (PD>4), for example, for the species *G. melonella, D. abietella* and *Scripophga incertulas*. In addition, the migratory and polyphaghous painted lady butterfly *V. cardui* also hosts two very distinct and novel iflaviruses; WUR31/32 and WUR20. The low number of clusters and the high phylogenetic diversity of the viruses suggests that the iflaviruses in these species have only been explored to a limited extent.

It is important to emphasize that the inferences that we make here on the host range of viruses strongly depend on the data that is available. Sequencing data is typically dominated by a few model organisms (Supplementary tables: Table S3), in which also potentially more viruses can be detected; in contrast, no or little data is available for most species impeding host range inferences. Furthermore, the infection status of the animals is generally unknown, and the detection of some viruses could point to contamination or viral infections in cell culture. Additionally, assigning the host of the virus from sequencing data can be difficult since samples generally consist of multiple organisms, including the target organism, its microbiome, potential parasites, and its gut content, all of which can also contain viruses. This is a general problem in virome and virus data mining studies (e.g., Zayed et al. 2022) and we aim to mitigate it here by focusing on iflaviruses, a virus group well known to infect insects, and particularly lepidopterans. Nevertheless, iflaviruses were also found to infect other organisms that could be present in insect data sets, such as plants and fungi (Coyle et al. 2024; Saqib et al. 2015). The discovery of iflaviruses in fungal pathogens lead to the proposition of new host assignments for viruses previously associated with ticks and *Drosophila* (Coyle et al. 2024). This report highlights the difficulty of host assignment based solely on metatranscriptomics data of animals and suggests additional analyses to verify the host assignment.

In this section, we detected similar iflaviruses across multiple lepidopteran species, suggesting that some members of the *Iflaviridae* family contain iflaviruses with a broad host range or they are capable of host jumps. An important predictor of virus susceptibility is the genetic distance between the novel host and the natural host of the virus (Longdon et al. 2014). By focusing only on Lepidoptera, an insect order that originated from nocturnal, herbivorous moth ancestors ∼100 million years ago in the Americas (Kawahara et al. 2023), we screen closely related hosts, which increases the probability of finding viruses with a broad host range. In addition, the main transmission route of iflaviruses is considered to be horizontal, which facilitates cross-species transmission and potential host jumps when different butterfly and moth species share habitats.

Besides iflaviruses, Lepidoptera can also be infected by other viruses (Ros et al. 2022), which can result in virus-virus interactions within a host individual as observed in several species. For example, in the Queensland fruit fly, *Bactrocera tryoni*, a negative association was found between the presence of an iflavirus and a sigmavirus (Sharpe et al. 2024). Furthermore, it was shown that the presence of one or both iflaviruses, SeIV1 and SeIV2, increased the pathogenicity of the baculovirus *Spodoptera exigua* multiple nucleopolyhedrovirus (SeMNPV, dsDNA virus) in *S. exigua* (Carballo et al. 2017). Our results indicate that from libraries of agricultural pests like *P. nuda, S. frugiperda*, and *E. obliqua*, new and known iflavirus genomes could be assembled. Specific baculoviruses, such as *Perina nuda* nucleopolyhedrovirus (Wang et al. 1999), *Spodoptera frugiperda* multiple nucleopolyhedrovirus (Barrera et al. 2011), and *Ectropis obliqua* nucleopolyhedrovirus, have been used as biological control agents against these pests. Whether the presence of iflaviruses affects the susceptibility and efficiency of baculoviruses in the above-mentioned species remains unclear. Our study highlights the presence of iflaviruses in agriculturally important Lepidoptera, which can be further explored for potential iflavirus-virus interactions and other cross-kingdom interactions, for example, involving viruses and bacteria.

### Genome evolution and recombination can be observed within an iflavirus species

In previous sections, we have shown that the assembly of public data largely expands the known iflavirus diversity, where also genomes of novel species can be reconstructed. Next, we used this data to zoom into the evolution of a particular virus species. We chose to analyze *Iflavirus betaspexiguae* in depth, since this is the species with the largest number of genomes: 274 complete genomes in addition to the two genomes at NCBI. We aligned the 98 non-identical protein sequences and analyzed the codon alignment. We detect a strong signal of recombination with 5 breakpoints (Figure 4). The alignment also shows a signal of diversifying selection when taking the partitions into account, with 40 sites under episodic diversifying selection (p<=0.1). However, when locating the positions under selection, we observed that only few positions show strong evidence (7 positions with p<0.01) and these are either between similar amino acids (BLOSUM80>=0) or can be found in only one genome, which makes it difficult to distinguish them from sequencing errors (Figure 4). Thus, although diversifying selection likely played a role in the evolutionary history of these genomes, the precise positions under selection cannot be located.

**Figure 4:**
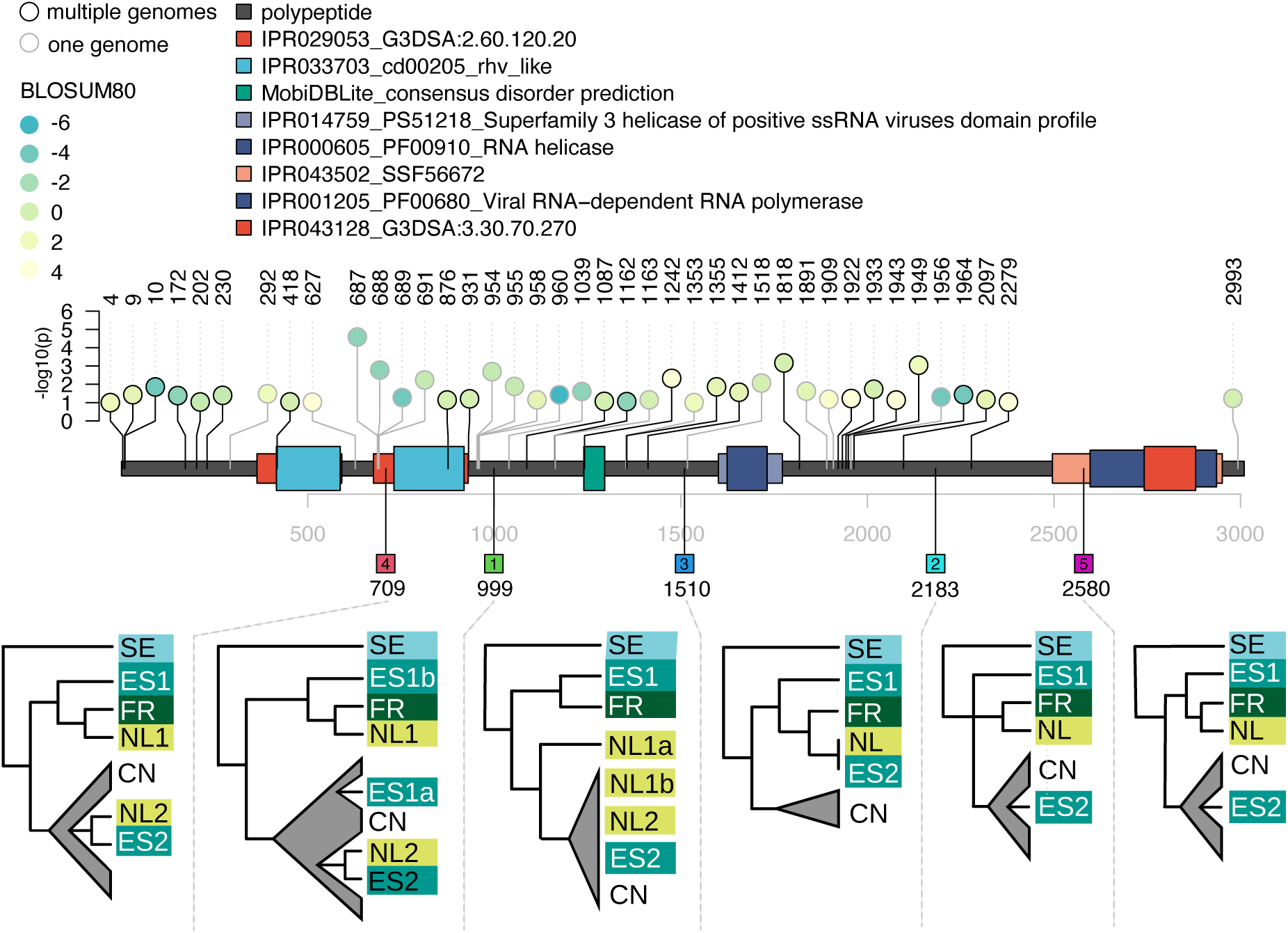
Genome evolution of *iflavirus betaspexiguae*. The polyprotein of YP_009010984.1 and the domains predicted by InterPro are shown. All domains with an IPR ID are shown and domains without an IPR ID are only shown if they do not overlap with other domains. Positions under diversifying selection are marked by circles on top of the genome, colored by how radical the amino acid change is (BLOSUM80 score, i.e., more negative corresponds to more radical) and if the change is in only one genome (gray) or multiple (black). The changes in one genome occur in AHX00961.1 (positions 687, 688, 689, 691, 954, 955, 958, 960), in SRR11822924_complete_1 belonging to ES2 (positions 292, 1353), and the remaining positions are all in different genomes. Recombination breakpoints are listed by squares on the bottom of the genome. Numbers within the square mark the order in which they were found, i.e., number 1 is the most supported. Phylogenies based on codon alignments are shown below the breakpoints. Groups are marked by the country which submitted the data set: SE (Sweden): SRR5464078_complete_1, SRR8269436_complete_1, ES1b: SRR1050534_complete_3, ES1a: SRR1050532_complete_2, SRR1050533_complete_3, ES1: ES1a and ES1b, ES2: SRR11822922_complete_1, SRR11822924_complete_1, ES (Spain): ES1 and ES2, FR (France): SRR13488425_complete_1, NL1a: SRR7415763_complete_2, SRR12002069_complete_1, SRR12002072_complete_1, NL1b: SRR7415760_complete_2, SRR7415766_complete_2, SRR12002067_complete_1, NL1: NL1a and NL1b, NL2: SRR7415761_complete_2, SRR7415762_complete_2, SRR7415767_complete_2, SRR7415768_complete_2, SRR7415771_complete_1, SRR12002063_complete_1, SRR12002066_complete_1, SRR12002073_complete_1, NL (Netherlands): NL1 and NL2, CN (mostly China): all the remaining genomes. Expanded phylogenies can be found in Figure S4.

A closer look at the phylogenies of the individual partitions shows that the positions of libraries from Spain and the Netherlands differ between the partitions (Figure 4). In particular, the samples from Spain group according to the two different transcriptomics experiments they were derived from (Park et al. 2014; Llopis-Giménez et al. 2020). Although the libraries from the Netherlands also originate from two different experiments (Breeschoten et al. 2019; Simon et al. 2021), we surprisingly observed that the grouping into NL1 and NL2 seems independent of the time of the experiment and the experimental condition (Supplementary tables: Table S6). The consensus sequences of NL1 and NL2 (Figure S5) are divergent in the structural region (310 single nucleotide polymorphisms (SNPs) in region 1 to 3750, 8.3% difference). However, the nonstructural region shows only two SNPs, at position 4230 (position 1410 in protein) and 6835 (position 2279 in protein). Position 1410 is a synonymous mutation, whereas position 2279 was also detected under selection due to a replacement between the chemically highly similar amino acids isoleucine and valine (Figure 4).

To test whether multiple variants of *Iflavirus betaspexiguae* coexisted over time in laboratory *Spodoptera exigua* populations from the Netherlands, we mapped the non-host reads to the consensus sequences of NL1 and NL2. Due to the high similarity in the nonstructural region, many reads map ambiguously in this region, which is observed by a mapping quality of 0. Nevertheless, the reads that map to a specific SNP can be used to distinguish the variants in the nonstructural region. We find that indeed NL1 and NL2 can be detected together in most of the samples (Figure S6). Additionally, investigating the coverages shows that many samples also support recombinants between NL1 and NL2 (Figure S6, Supplementary tables: Table S6), with at least two recombination breakpoints: one between genome positions 3750 and 4230 and one between genome positions 4230 and 6835. Due to the high similarity in these regions, the breakpoints cannot be narrowed down further. Interestingly, the first breakpoint identified lies between protein positions 1250 and 1410,placing it at the boundary between the structural and non-structural regions (Figure 4). These results demonstrate that multiple variants of *Iflavirus betaspexiguae* can be found in laboratory samples, where the high genomic similarity between them allows them to engage in recurrent recombination.

Here we showed that a vast genomic diversity exists within an iflavirus species. This diversity was previously underestimated due to two steps in the analysis. First, a clustering step is often applied to reduce diversity prior to phylogenetic analysis, which masks the complex evolutionary dynamics that occur within virus species. Second, assembly methods can only reconstruct the consensus sequence when a high diversity of variants is present in the samples. This can lead to fragmented assemblies and to the reconstruction of consensus sequences that do not correspond to a naturally existing virus genome. Here we observed within-sample diversity for selected samples; in subsequent analyses, additional methods to study genetic diversity within samples could be applied to decipher closely related strains (e.g., Fuhrmann et al. 2024). Furthermore, we find that for some samples, like the ones from Spain, the different virus variants are associated with different transcriptomics experiments, suggesting that different insect populations have been used for the experiments. For the samples from the Netherlands, we showed that different closely related variants co-exist and evolve by mutations and recombination within experimental time scales. Since viruses evolve rapidly, it was also suggested to use them as fast-changing markers to study the ecology of their hosts, for example for mosquitos (Hollingsworth et al. 2023). Here we detected a high diversity of iflaviruses in Lepidoptera and demonstrate their potential to evolve rapidly, which makes them also excellent candidates to study the evolution and ecology of RNA viruses and their hosts.

## Conclusion

Here we presented a data mining study based on publicly available sequencing data from Lepidoptera transcriptomes. Our analysis expanded iflavirus diversity, by finding novel viruses and by increasing diversity within known virus species. This allowed us to expand the host range of iflaviruses across lepidopteran species, where similar iflaviruses could be found in different host species and genera and where divergent iflaviruses were present in one host species. By studying the diversity of one virus species, we reconstructed its evolutionary dynamics and found pervasive signatures of recombination. Additionally, we developed a pipeline to automatically reconstruct viruses from public transcriptome data, which includes automatic download from SRA, mapping, assembly, and virus similarity search; this pipeline can be applied to other virus families and hosts of interest. This study serves as a proof of concept for reconstructing intra-family viral diversity from public sequencing data, providing a basis for studying virus evolution.

## Methods

### Data gathering using Serratus

The Serratus database (version from 3 April 2023) was used to find libraries from the SRA database at NCBI that contain reads mapping against the RdRp consensus sequence of one of the families of the viral order *Picornavirales* according to the International Committee on Taxonomy of Viruses (ICTV; *Picornaviridae, Secoviridae, Iflaviridae, Marnaviridae, Dicistroviridae*) (Valles et al. 2017). In addition, we added 30 publicly available libraries from one *Spodoptera exigua* population at Wageningen University to the list (SRR7415760 - SRR7415771 and SRR12002056 - SRR12002074). Serratus detected signatures of iflaviruses in 15 of the 30 libraries (Supplementary tables: Table S1). To filter for hosts of the order Lepidoptera, we filtered the European Nucleotide Archive (ENA) for sequencing runs that have species belonging to the lepidopteran order as a model organism (filter ‘tax_tree(7088)’, 29 March 2023). The overlap of the lists of accessions from Serratus and from ENA yielded a total of 5,035 eligible libraries (Figure 1A).

### Virus genome reconstruction

The workflow management system Snakemake (v5.26.0) (Mölder et al. 2021) was used to develop an automated pipeline in combination with the software package manager Conda (v22.11.1). Snakemake user-defined resources were used to prevent the pipeline from exceeding the storage limit and the number of allowed application programming interface (API) calls to NCBI. The pipeline code, a conda environment file, and test data to run the pipeline are available in a GitHub repository (https://github.com/annecmg/VirusRePublic).

#### Raw data download

To include only paired-end (PE) RNA-Seq libraries, FFQ (v0.3.0) was used to check whether two fasta files were stored on the NCBI FTP server (Gálvez-Merchán et al. 2023). Of the 5,035 eligible libraries, 4,104 (82%) were of paired-end library layout and included in the analysis.

The compressed reads data (.sra format) of the included accessions was downloaded directly from the Amazon Web Services (AWS) mirror of the SRA using prefetch (v2.9.1) (SRA Toolkit Development Team 2018). We observed that AWS limits the download speed of jobs when a high amount of bandwidth is requested for an extended period, which can drastically increase the time that is needed to download the reads. To overcome this issue, each download was allowed to run for 20 minutes. If not successful, the download of the compressed data was automatically restarted, which typically increased bandwidth and let to faster completion. Subsequently, the data was decompressed into two FASTQ files per accession, using fasterq-dump (v2.9.1) (SRA Toolkit Development Team 2018).

#### Host genome download

The metadata of all the PE SRA libraries was downloaded using a combination of esearch and efetch included in the Entrez Programming Utilities package (v16.2) (Rédei 2008). Using this package, all the unique NCBI taxonomic identifiers (TaxIDs) of the organisms included in the libraries were extracted from the metadata. Subsequently, the NCBI datasets command-line tool (v14.6.0) was used to determine for which of these TaxIDs a genome assembly was available on the NCBI server (O’Leary et al. 2024). Host genome assemblies classified as the reference genome were preferred and otherwise the assembly with the highest contig N50 was downloaded. Host genome assemblies were downloaded using a recently developed method to download large amounts of genomic data included in the NCBI datasets command-line tools. In short, dehydrated data packages (< 5 KB in size), containing metadata and the location of the full data, were downloaded for each genome. These packages were then unzipped and rehydrated to obtain the genome sequences. Host genomes were indexed using HISAT2 (v2.2.1) (Kim et al. 2019).

This yielded a total of 329 unique hosts TaxIDs of which 31 had a RefSeq genome assembly available (comprising 2,400 libraries), 119 with a GenBank assembly (comprising 1,144 libraries) and 179 had no genome assembly (comprising 458 libraries). For 102 SRA libraries, the host organism could not be determined using the metadata files (e.g., due to the TaxID being specified in the wrong field, or the TaxID being defined as multiple organisms) and these were excluded. The 4,002 accessions where the host was determined were included in the analysis (Supplementary tables: Table S1, Figure 1A).

#### Read processing

We used fastp (v0.23.2, --*dedup, --dup_calc_accuracy 6, --thread 3*) to perform adapter and quality trimming of the reads, as it can automatically detect adapter sequences and trim the reads accordingly (Chen et al. 2018). For SRA libraries where a host genome was available, read mapping was performed using HISAT2 (*-t, -p 4, --no-temp-splicesites*) and the HISAT2 SAM output was converted to BAM format (Samtools v1.16.1, *view --threads 4, -Sbh)* (Danecek et al. 2021). Unmapped reads were extracted using Samtools (*view --threads 4, -h -b, -f 4*), and converted to compressed PE FASTQ format (fastq.gz) using Samtools (*fastq --threads 4 -N*).

#### Genome assembly

rnaviralSPAdes v3.15.5 was used for assembly. We aimed to provide an appropriate K-mer list to the assembler, starting with the original K-mer list [33, 55, 77, 99, 127]. As the read length varies between SRA libraries, the seqkit stats option was used to determine the average read length (after trimming) for each library (v2.3.1) (Shen et al. 2016). Then, the highest K-mer in the list was set to the K-mer from the original list that is closest to 75% of the read length of a library. For example, if the average read length is 100 nucleotides (nt), [33, 55, 77] was used. Additionally, the snakemake pipeline catches assemblies that fail with exit code 21 (used by SPAdes to identify instances with read length < K-mer size), removes the highest K-mer from the list and restarts the assembly. The snakemake pipeline also handles the rare occasion when SPAdes exits with code 255, which happens when an SRA run is classified as PE data and two files are available, but the data in reality consists of SE reads spread over two separate files. Then an empty dummy contig file is created.

#### Viral protein identification

To detect assembled contigs with similarity to *Picornavirales* contigs, the sequence aligner DIAMOND was used (v2.0.15.153) (Buchfink et al. 2021). First, a database of all known *Picornavirales* proteins was created (25 April 2023), by directly downloading the fasta sequences assigned to *Picornavirales* from the NCBI protein database, using the search term “Picornavirales”[Organism], resulting in 206,790 included protein sequences. Assembled contigs were aligned against this protein database (*diamond blastx, -p 2, -f 6*). When DIAMOND encountered an empty contig file, a dummy output file was created, and the specific library was logged as corrupted. We filtered the DIAMOND output for hits to the different *Picornavirales* families. For non-*Iflaviridae*, significant contigs (e-value threshold 0.001) with a length of at least 5000 nt are considered as putatively viral.

The assembly was performed using the unmapped reads when a host genome was available and for all the reads otherwise. To confirm that there was no quality difference between *de novo* assembly and assembly of unmapped reads after host genome mapping, a test set of 100 libraries that had a host genome assembly available, was analysed using both methods. For these 100 libraries, both DIAMOND alignment files were compared and showed the same targets with almost identical statistics. Hence, we conclude that the host genome mapping only sped up the running time of the workflow but did not influence the quality of the results.

#### Determining completeness for iflavirus contigs

For *Iflaviridae*, we aimed to reconstruct the complete polyprotein. First, a contig must have a best hit with the following criteria: (i) the product of the alignment length and identity percentage (pAID) is over 30,000 and (ii) the best hit belongs to *Iflaviridae*. The pAID cutoff was used to filter for alignment length and identity percentage at once. For example, some contigs showed alignments with high sequence similarity against the viral RdRp only, which was filtered due to their short length. Next, if a contig matched the criteria and was between 8,000-12,000 nt, the contig was classified as being a ‘complete’ *Iflaviridae* genome. If the length of the contig was less than 8,000 nt and there were no other contigs passing the criteria above, the sequence was labelled as an ‘incomplete genome’. If there were multiple contigs hitting the same target with the criteria above that were shorter than 8,000 nt in length, the sequences were classified as ‘fragmented genome’. To reconstruct multiple iflavirus genomes in one library, all the alignments to contigs that contribute to the reconstructed genome were removed and the above procedure was repeated until no more genomes could be reconstructed. Complete genomes were translated to proteins using EMBOSS *(-minsize 6000 -find 1*).

#### Patching fragmented genomes

A total of 384 genomes were classified as being fragmented, which we attempted to ‘patch’ into a protein sequence. The contigs that corresponded to the fragmented genome were translated into a protein sequence using EMBOSS getorf (*-minsize 700, -auto*). The protein sequences were aligned against the local DIAMOND protein database using the blastp option (*blastp, -p 2, -f 6, --quiet*) and sorted on alignment length in descending order. A template sequence was created containing as many ‘X’ characters as the length of the best protein target. Then the template sequence was modified by replacing the region between target start and end positions provided in the DIAMOND blastp file with the subject protein sequence, where only alignments against the best protein target are considered. If the aligned region of one or multiple sequences overlapped, the amino acid sequence of the first contig was preferred. The final patched sequence was required to contain at least 2,150 non-X characters, which was chosen to be 80% of the length of the shortest, best target that was used as template sequence (AAQ17044, 2,674 aa in length). After the patching process, 240 patched genomes (63%) passed this cut-off.

### Evolutionary analysis

#### Iflavirus proteins from NCBI

Iflavirus proteins were downloaded from NCBI virus (https://www.ncbi.nlm.nih.gov/labs/virus) on 22 August 2023. Restricting to Virus “Iflaviridae, taxid:699189” and Nucleotide Completeness “complete” yielded 322 protein sequences with a length of at least 2,000 aa. From this list, 35 GenBank proteins with an identical assembly in RefSeq were removed. Additionally, six accessions were included (QID77674.1, QID77679.1, QHI42120.1, QJB76116.1, QQN90112.1, WAX26121.1), where the taxonomy was not annotated but that showed high identity when the reconstructed genomes were searched against nr with blast (see below). The resulting list of 293 accessions from NCBI contains 45 RefSeq and 248 GenBank accessions (Supplementary tables: Table S2).

#### Redundancy clustering

We clustered the reconstructed iflaviruses with the ones from NCBI based on pairwise blastp with identity of 99% and alignment coverage of 95% on at least one sequence. Since the patched genomes contained X characters which are considered as mismatches in blast, X characters have been removed for this step. Consequently, identity is calculated as the number of identical matches divided by the number of identical matches plus the number of mismatches, i.e., gaps are ignored for the identity calculation. As cluster representative, the following priority was considered: (i) NCBI genomes with RefSeq accession, (ii) NCBI genome with GenBank accession, (iii) complete reconstructed genome, (iv) patched reconstructed genome and the longest genome in a category was selected.

To confirm that the reconstructed genomes indeed belong to *Iflaviridae*, clusters that did not contain an NCBI genome were searched with BLASTp 2.14.0+ against the nr database (downloaded 9 March 2023). The results were parsed with TaxonKit to obtain the lineage of the best hit (Shen & Ren 2021; Camacho et al. 2009). Manual inspection of the best hits that did not belong to *Iflaviridae* showed that these sequences were generally classified as *Riboviria* and a detailed taxonomic classification was missing. These sequences have been added to the included NCBI genomes (see above) and the clustering was repeated.

#### Phylogenetic analysis

Two different data sets were obtained. One for all the cluster representatives (termed “all-host data set”) and one with only the cluster representatives from lepidopteran hosts (termed “Lepidoptera data set”). For each of the data sets, protein sequences were aligned using MAFFT v7.490 with the linsi algorithm (Katoh & Standley 2013). The alignment was trimmed with TrimAl (-gt 0.6 -w 3) (Capella-Gutiérrez et al. 2009). Next, a phylogenetic tree was reconstructed from the trimmed alignment with IQ-TREE v2.2.6 (-B 1000) and the best selected model (Minh et al. 2020; Kalyaanamoorthy et al. 2017). Phylogenetic trees were visualized using iTOL as midpoint rooted (Letunic & Bork 2024).

Based on the iflavirus phylogeny for the Lepidoptera data set, the phylogenetic diversity (PD) is calculated for each host species. To calculate PD for a host species, we use ETE 3 to prune the tree to all nodes where the cluster contains at least one sequence from this species while preserving the branch lengths (Huerta-Cepas et al. 2016). PD is then the sum of branch lengths of the pruned tree.

#### Pairwise distance calculation

According to the ICTV criteria (https://ictv.global/report/chapter/iflaviridae/iflaviridae/iflavirus), iflaviruses are assigned to the same species if they have at least 90% amino acid identity in their capsid proteins (Valles et al. 2017). Based on the iflavirus multiple sequence alignment for lepidopteran hosts and the annotation provided in the NCBI genomes, an alignment position was determined, which marks the end of the capsid proteins (alignments can be found at https://github.com/annecmg/VirusRePublic/tree/main/Data_Paper_Lepidoptera/alignments). The N-terminal alignment until that position was considered as the alignment of capsid proteins and pairwise distances are calculated from the multiple sequence alignment using EMBOSS distmat and uncorrected distances (Rice et al. 2011). Pairwise identity was calculated as 1-distance.

#### Iflavirus species assignment

For each cluster representative that is a complete genome, we looked for the highest pairwise identity (see previous section) to an NCBI genome with an assigned species. If that identity was at least 90%, the complete genome was assigned to the same species as the identified NCBI genome. Otherwise, if any NCBI genome with pairwise identity at most 90% could be found, the complete genome was considered as a known genome; otherwise, it was considered as a “novel genome”. The assignment for the cluster representative was transferred to all cluster members.

#### Calculation of lowest common ancestor (LCA)

The lineage for each host species in the data set was obtained from NCBI and genus and family were extracted using ETE 3. The LCA for a pair of genomes was determined in a stepwise manner: if the genomes have the same host species, its LCA was assigned to “Species”; otherwise, if the genomes share the host genus, its LCA was assigned to “Genus”, otherwise, if the genomes share the host family, its LCA was assigned to “Family”, otherwise, the LCA was assigned to “Order” (i.e., Lepidoptera). Accordingly, the LCA of a cluster is the highest rank among the LCAs of all cluster members.

#### Domain annotation

Cluster representatives were annotated with InterProScan v5.68-100.0 (Paysan-Lafosse et al. 2023).

#### Analysis of Iflavirus betaspexiguae

We included the two genomes from NCBI and the 274 complete genomes assigned to *I. betaspexiguae* in an in-depth evolutionary analysis of this virus species. We deduplicated these genomes, by removing sequences with 100% identity over 100% of the length to another sequence, i.e., an identical or longer sequence was still included, resulting in 98 genomes. Protein sequences were aligned using MAFFT v7.490 with the linsi algorithm (Katoh & Standley 2013) and subsequently, a codon alignment was generated using PAL2NAL (Suyama et al. 2006). Recombination in the codon alignment was assessed using GARD in the HyPhy package (Kosakovsky Pond et al. 2006). Diversifying selection was detected using MEME in the HyPhy package based on the partitioned alignment (Murrell et al. 2012). Phylogenies of partitions detected by GARD were re-estimated with IQ-TREE v2.2.6 (-B 1000) and the best selected model (Minh et al. 2020; Kalyaanamoorthy et al. 2017). Consensus sequences were reconstructed with EMBOSS cons (Rice et al. 2011). For some libraries, unmapped reads (regarding the host genome) were mapped to virus genomes using bwa mem (Li 2013). Mappings were visualized in R using the bioconductor packages GenomicAlignments and Rsamtools (Lawrence et al. 2013; Morgan et al. 2024). Plots were produced with ggplot2 (Wickham 2016).

## Data availability

Nucleotide sequence data of the 65 cluster representatives originating from the reconstructed complete genomes (WUR1-WUR65) are available in the Third Party Annotation Section of the DDBJ/ENA/GenBank databases under the accession numbers TPA: BK068472-BK068536 (BioProject PRJNA1137245). The sequences with similarity to other *Picornavirales*, the complete and patched iflavirus genomes, and the described alignments are available in a GitHub repository (https://github.com/annecmg/VirusRePublic/tree/main/Data_Paper_Lepidoptera). The pipeline code, a conda environment file, and test data to run the pipeline are available in a GitHub repository (https://github.com/annecmg/VirusRePublic).

## Acknowledgements

We would like to thank Thomas de Bruijn, Dimitris Karapliafis (DK), Jun Liu, and Laura Pantino Medina for comments on the manuscript. We also appreciate the development of earlier versions of the virus detection pipeline by Matthijs Kon and Matthijs Pon and testing of the final pipeline by Arjan Draisma and DK.

## Author Contributions

Devin van Valkengoed: Methodology, Analysis, Writing – original draft, Writing – review & editing. Astrid Bryon: Conceptualization, Writing – original draft, Writing – review & editing. Vera I.D. Ros: Conceptualization, Writing – review & editing. Anne Kupczok: Conceptualization, Methodology, Analysis, Writing – original draft, Writing – review & editing.

## Supplementary figures

**Figure S1:**
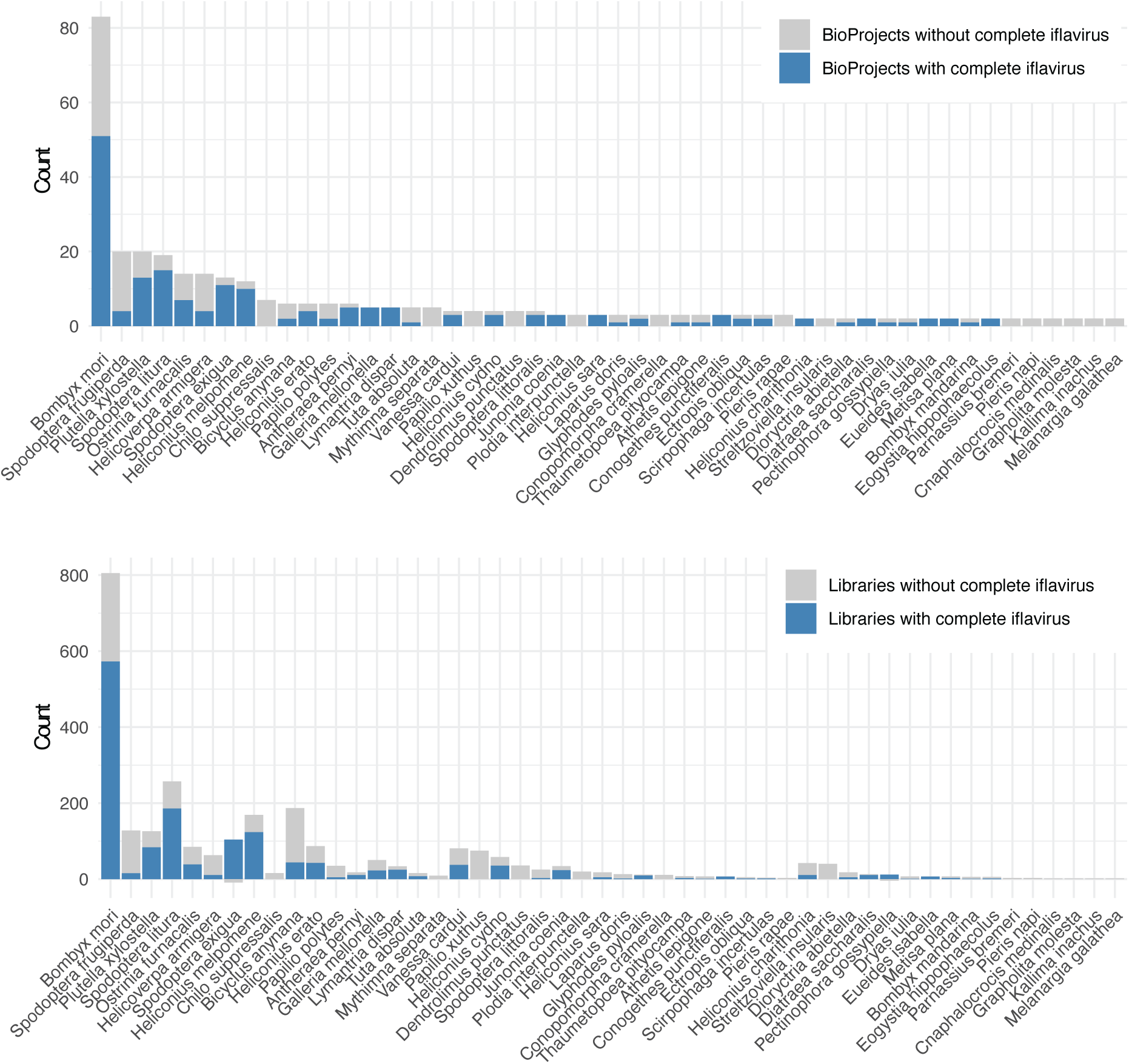
Number of BioProjects (top) and libraries (bottom) where iflaviruses where detected. Note that complete genomes were only reconstructed for some of them. Only species with at least two included BioProjects were included in the plot; full data can be found in Supplementary tables: Table S3.

**Figure S2:**
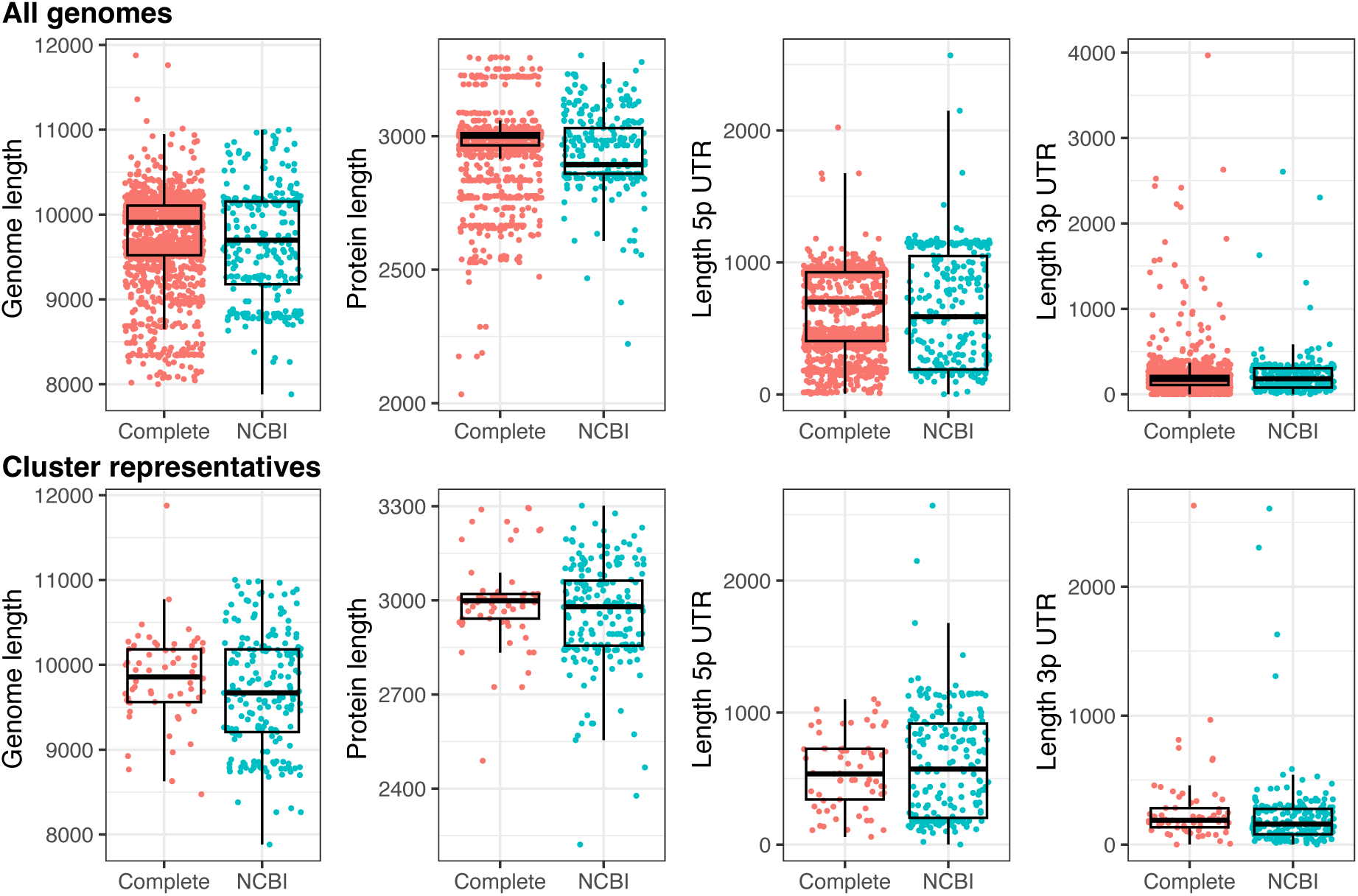
Genome statistics for reconstructed complete genomes and genomes from NCBI.

**Figure S3:**
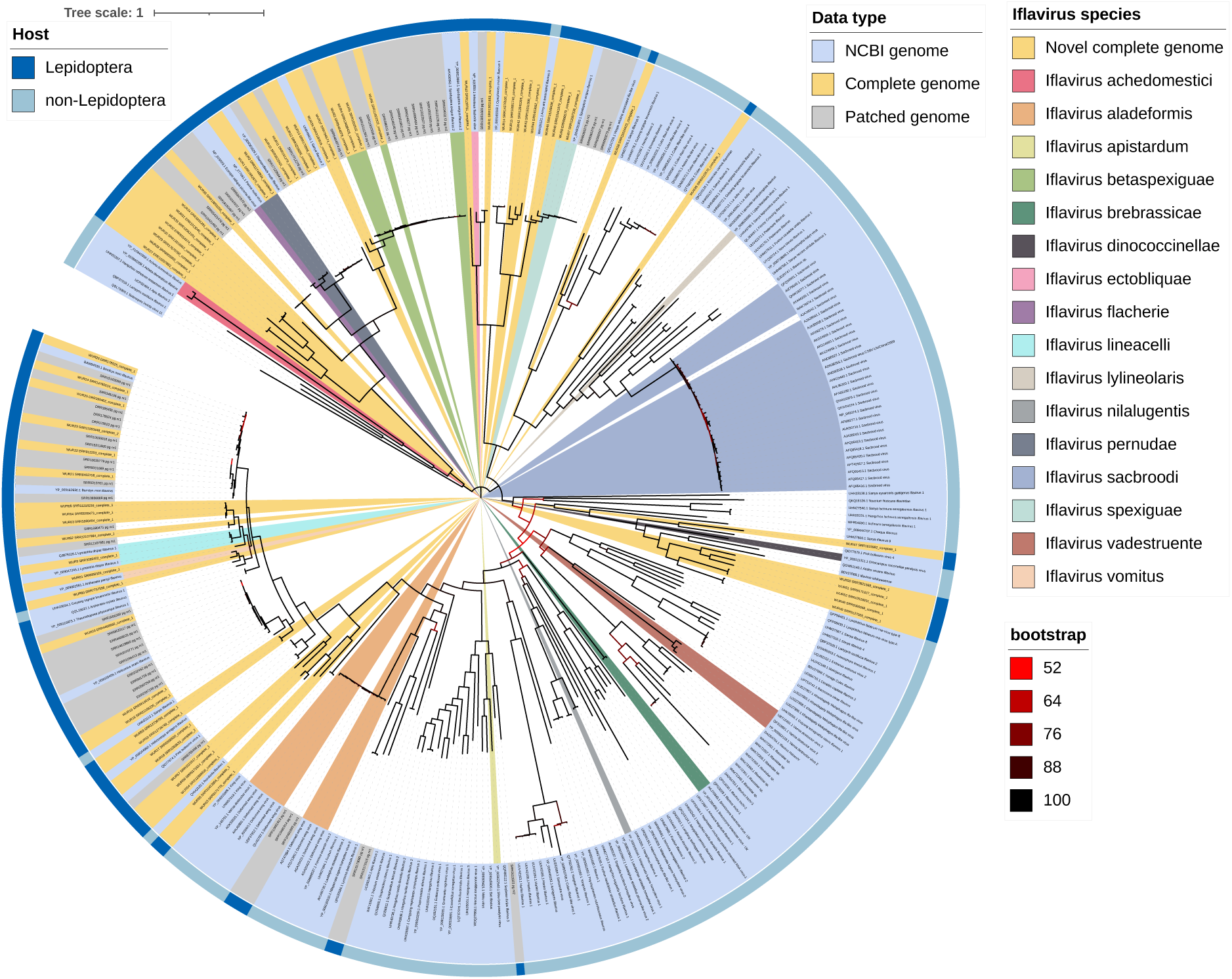
Iflavirus phylogeny for the all-host data set. The outer ring denotes the host, taxa name is colored by data type, clades are colored by iflavirus species, and branches are colored by bootstrap value. A fully annotated phylogeny can be found at https://itol.embl.de/tree/31201033175461721724737.

**Figure S4:**
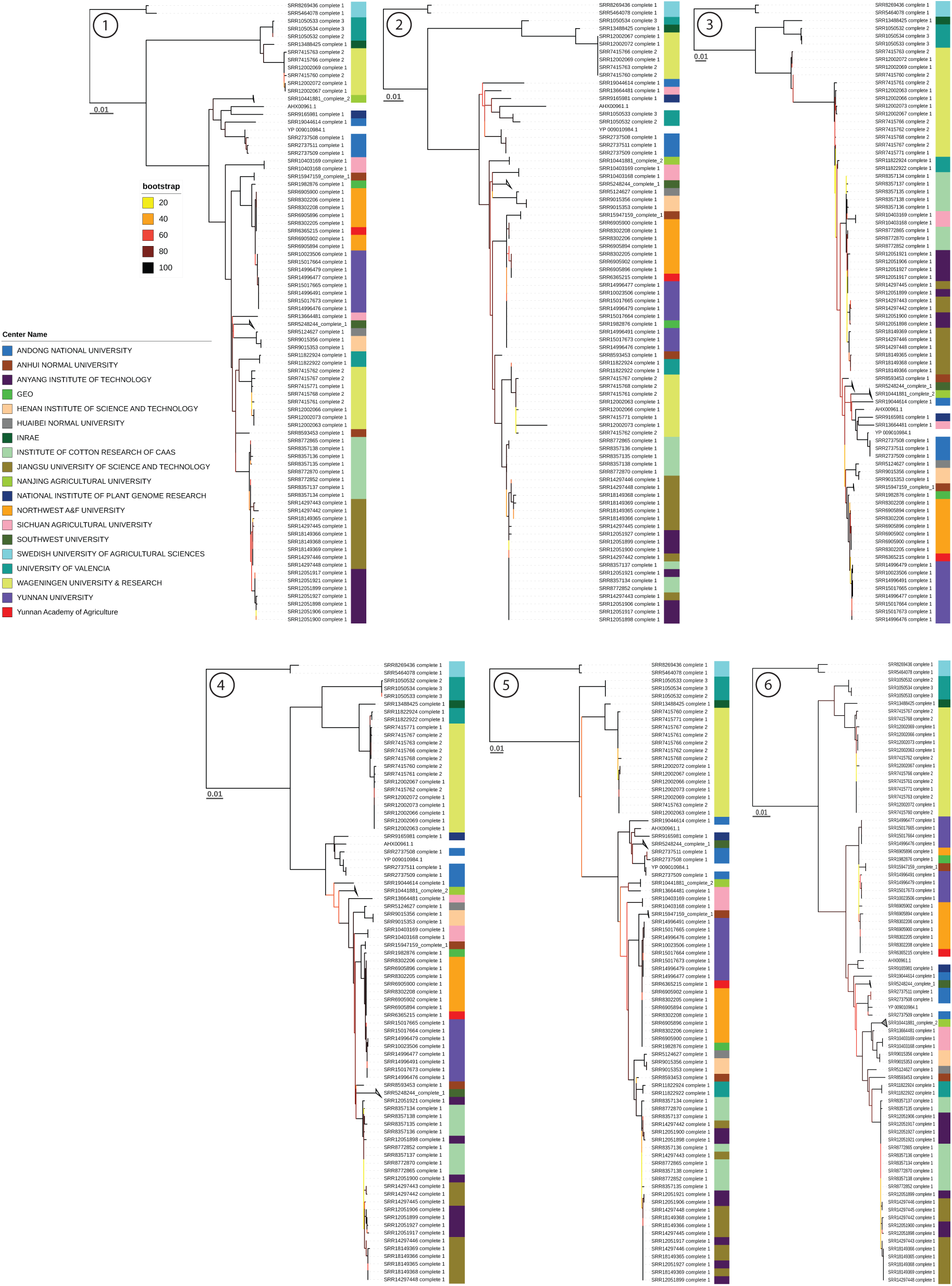
*Iflavirus betaspexiguae* trees per partition (given as number in circle). Groups with at least 3 genomes that are present in all trees are collapsed and labelled by one representative: 14 genomes from Southwest University are collapsed and labelled SRR5248244_complete_1, 4 genomes from Nanjing Agricultural University are collapsed and labelled SRR10441881_complete_2, 3 genomes from Anhui Normal University are collapsed and labelled SRR15947159_complete_1.

**Figure S5:**
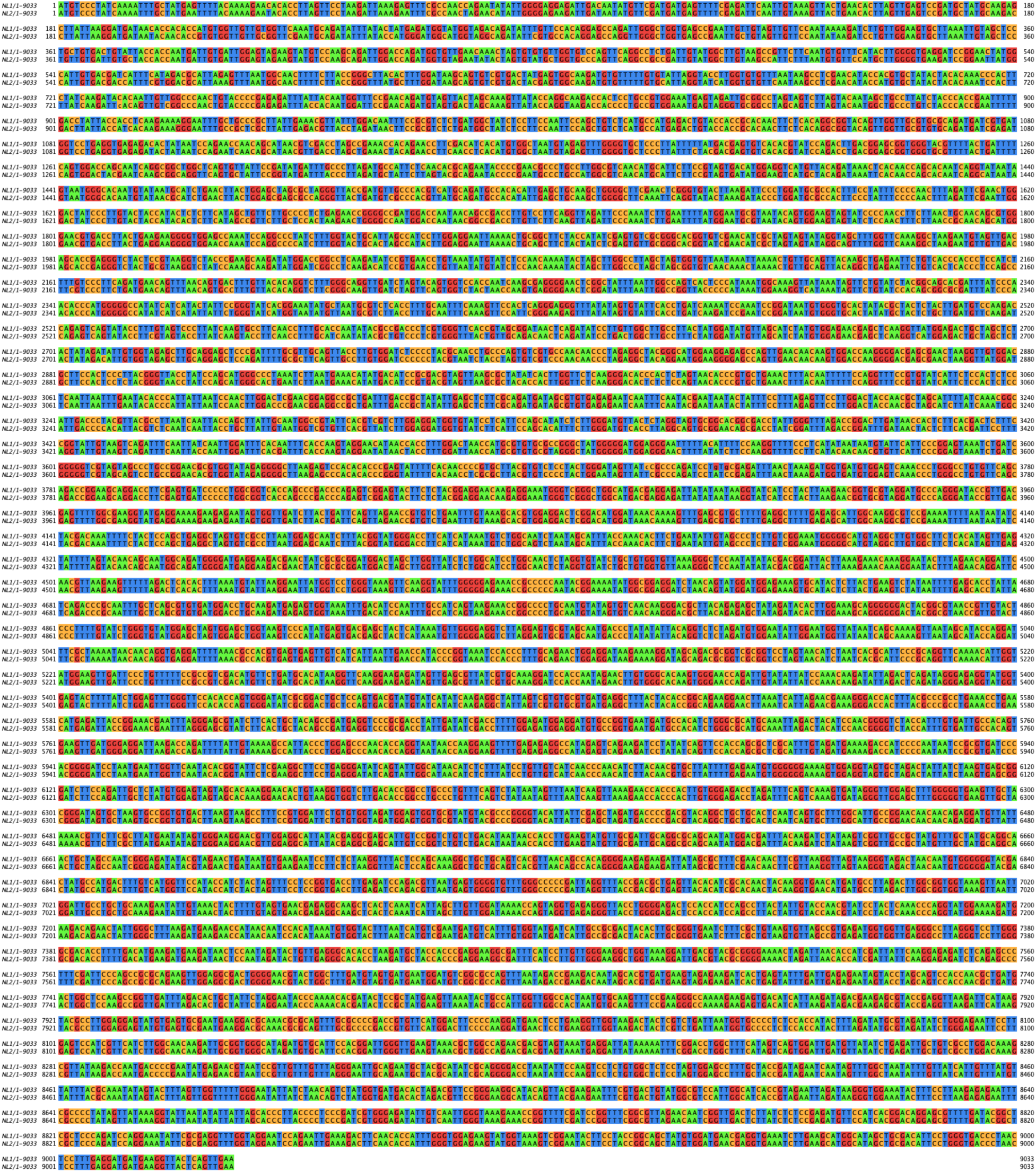
Consensus sequences of NL1 and NL2 displayed as an alignment.

**Figure S6:**
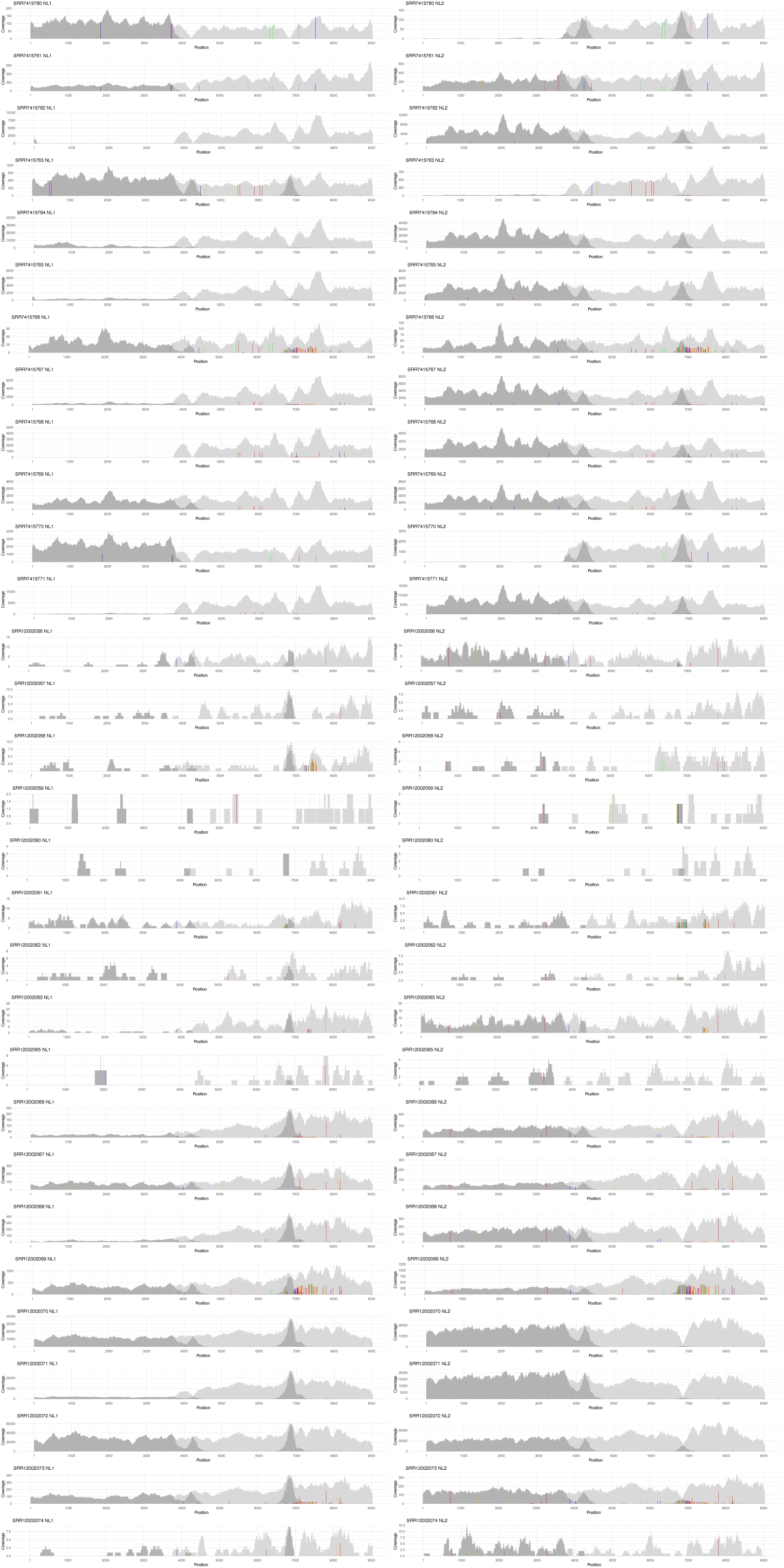
Mapping of samples from the Netherlands against consensus of NL1 and NL2 (Figure S5). The reads were mapped against both sequences at the same time (i.e., competitively). The coverage along the genomes is displayed in light gray for all mapped reads and in dark gray for mapped reads with mapping quality > 0 (i.e., uniquely mapped reads). Since the non-structural region is highly similar, most of the mapped reads in this area cannot be mapped uniquely. Nevertheless, the non-structural region differs by two SNPs (positions 4230 and 6835) and the presence of NL1 or NL2 can be seen by dark peaks at these two locations. Single nucleotide variants (SNVs) that differ from the reference and are supported by at least 2 reads are displayed as colored bars (A – green, C – blue, G – orange, T – red). Different versions and recombinants can be observed (see also Supplementary tables: Table S5): For example, SRR7415760 maps mainly to NL1 in the structural region and to NL2 in the non-structural region. SRR7415763 contains a variant of NL1, with few SNVs, and also NL2 at lower frequency. SRR7415766 contains both NL1 and NL2 and an additional variant in the non-structural region that is characterized by many SNVs. SRR7415768 contains mainly NL2. SRR12002066 contains both NL1 and NL2 but shows mainly the NL2 version at position 6835 indicating a recombination breakpoint between the two SNPs.

## Legends for Supplementary tables

**Table S1: Included SRA libraries**. Metadata provided by SRA, indication for which viral families a library was detected by Serratus, and the numbers of reconstructed genomes.

**Table S2: Included iflavirus polyproteins from NCBI**. The organism and host columns originate from the NCBI metadata. Host species and Host order have been inferred from the metadata information or by manually looking up the NCBI entry and publication.

**Table S3: Included libraries and detected and reconstructed iflaviruses per host species.** For hosts with complete iflavirus genomes, the number of iflavirus clusters and the phylogenetic diversity is also provided.

**Table S4: Presence of InterPro domains in the cluster representatives.** InterPro accessions are only listed if they occur in the Lepidoptera data set. Accessions found in nearly all the NCBI genomes from the Lepidotera data set are marked in bold. The NCBI genome from the Lepidoptera data set lacking IPR014759 (Helicase, superfamily 3, single-stranded RNA virus) is UHR49801.1 Sanya iflavirus 2. The complete genome lacking the polymerase (IPR043502, IPR001205, and IPR043128) is WUR9 SRR3089402_complete_1.

**Table S5: Number of genomes detected for each iflavirus species.** “Novel” marks genomes where, according to ICTV criteria, no related genome at species level is known. Note that species assignment was not attempted for clusters containing only patched genomes, resulting in an overestimation of “novel” patched genomes. Of the 93 “novel” clusters, 39 contain a complete genome and can be considered novel species.

**Table S6: Assembly and mapping results for samples from the Netherlands.** Coverage across different regions of NL1 and NL2 (Figure S5) is calculated using samtools depth with a minimum mapping quality of 1.

